# Green microalga *Trebouxia* sp. produces strigolactone-related compounds

**DOI:** 10.1101/195883

**Authors:** I. Smýkalová, M. Ludvίková, E. Ondráčková, B. Klejdus, S. Bonhomme, O. Kronusová, A. Soukup, M. Rozmoš, B. Guzow-Krzemińska, R. Matúšová

**Author notes:** **Highlight** In lichenized alga *Trebouxia arboricola* there are produced SLs-related compounds inducing germination of parasitic weed *Phelipanche aegyptiaca; T. arboricola* stimulate growth of *Arabidopsis* roots and pea shoots; expression of SL-related genes in *P. patens*; carlactone detected in *T. arboricola*. **Abbreviations** PpCCD7: *Physcomitrella patens* CCD7 SL: Strigolactone GR24: Synthetic strigolactone analog DESI-MSI: Mass Spectrometry Imaging HPLC: High Performance Liquid Chromatography MRM-LC-MS/MS: Monitoring-liquid Chromatograhy-tandem Mass Spectroscopy.

## Abstract

Different algal species that may have germination inducing activity of holoparasitic broomrape weeds *Phelipanche aegyptiaca* and *P. ramosa* seeds were screened through germination bioassay. Green alga produce SL-related compounds. Applied extracts of biomass obtained from the culture of green alga *Trebouxia arboricola* increased seeds germination of both parasites. An optimatization of the alga extraction led to an increase of *P. aegyptiaca* germination. Exhausted medium also contained SL-related compounds. The crude extract stimulated the roots length of *Arabidopsis thaliana* tested *in vitro*. A similar effect had the algae and GR24 aplications on expression levels of the SL-related genes in *Physcomitrella patens*. The novel analytical method DESI-MSI detected production of carlactone in the algae. The *Trebouxia* sp. culture applications in pot experiments had positive effect on growth characteristics of pea plants.

## Introduction

The first report about the production of some compounds stimulating seed germination of obligatory root parasites: whichweed *Striga lutea*, broomrapes (*Orobanche* and *Phelipanche* spp.), and *Alectra* spp., has already been published in 1966 (Cook *et al.,* 1972). These compounds were recognized as growth stimulating factors exuded by host plant roots, later named strigolactones (Akiyama and Hayashi, 2006). The strigolactones (SLs) are the carotenoid-derived terpenoid lactone compounds produced in plant root tissues at low concentrations (10^-13^ M). The SLs are derived from carotenoids (Matusova *et al.,* 2005), a group of physiologically important compounds with conjugated double bond system synthetized by all photosynthetic organisms. SLs appearred in other part of plants such as hypocotyls, stems and leaves (Yoneyama *et al.,* 2007). Strigolactones were recognized as novel plant growth hormones acting similary to auxins, controlling shoot branching (Gomez-Roland *et al.*, 2008, Umehara *et al.*, 2008). The primary functions of SLs are inhibition of outgrowth of axillary buds into branches, stimulation of root branching in the plant (Brewer *et al.,* 2013), and they have a great impact on plant architecture which corresponds with long-distance transport of SLs (Kohlen *et al.,* 2011). The second important role is the stimulation of spore germination and hyphal branching of arbuscular mycorrhizal (AM) fungi by exudation of trace amounts of unstable SLs from roots of plants into rhizosphere (Besserer *et al.,* 2006, Bouwmeester *et al.*, 2007). GR24, a synthetic analog of strigolactones is the most extensively used SL analog in strigolactone research. Both production and exudation of SLs is increased in plants under stress conditions as nutrient deficiency, namely phosphorus limitation (Yoneyama *et al.,* 2007). These facts suggest a great problem mainly in Africa, and possible invasive expansion of parasitic plants into Mediterranean areas of Europe (Chauhan and Mahajan, 2014). The current knowledge about strigolactones implies no sufficient biological nor economic resolutions to solving this problem.

For the determination and study of structures of natural SLs in plants, the hydroponic system (cultivation of seedlings in liquid medium) is prefered. The root exudates or root extracts containing SLs are collected and concentrated (SPE C18 Cartridge) (Yoneyama *et al.,* 2012). The extracts or exudates are fractioned by HPLC and endogenous SLs are identified by multiple reaction MRM-LC-MS/MS. The disadvantage of SLs is inherently their unstability in aqueous solutions and also in solvents such as metanol. The biological activity of SLs strongly depends on stereochemistry (Zwanenburg and Pospíšil, 2013). The SL signaling requires the hormone-dependent interaction of α/β hydrolase DWARF14/D14 producing an intermediate molecule CLIM (De Saint Germain *et al.,* 2016, Yao et al., 2016), F-box of protein MAX2/D3 (Hamiaux *et al.,* 2012, De Saint Germain *et al.,* 2013), transcription regulator with proteolytic function SMXL7/D53 (Bennet *et al.*, 2016) and an uknown transcription factor (Li *et al.,* 2017). Several mutants of *Arabidopsis thaliana*, *Pisum sativum*, *Oryza sativa*, *Zea mays*, *Petunia hybrida* and *Physcomitrella patents* were used to identify SLs biosynthesis and signal transduction.

Carlactone is an endogenous biosynthetic precursor for strigolactones (Alder *et al.,* 2012, Seto *et al.,* 2014). In plant kingdom, there are three families of naturally occuring SLs, strigol-type, orobanchol type, and other SLs-like compounds, which are derived from the molecule of carlactone. The carlactone is easy oxidated by the enzyme MAX1, a cytochrome P450, to produce carlactonic acid and methyl carlactonoate (Abe *et al.,* 2014). This confirmed that carlactone is a precursor of wide variety of SLs-like molecules.

The soil algal biomass varies from 0 to 10^8^ cells g^-1^ soil (dw), mean value is calculated as 10 kg ha^-1^ and the highest mean abundance of algal cells occurs in the 0 - 2 cm soil layer. The main characteristics of soil algae are an excretion of organic acids that increase P-availability and P-uptake, provision of nitrogen by biological nitrogen fixation, increase in soil organic matter, production and release of bioactive extracellular substances that may influence plant growth and biosynthesis of plant growth regulators, crust formation, biofertilizers and biopesticides, stabilization of soil aggregation by extracelular polysacharides of soil aggregate and accumulation of metal ions present in their environment. Blue-green algae, especially the nitrogen-fixing cyanobacteria *Nostoc muscorum, N. calcicola, N. piscinale, Anabena sp., A. oryzae, Microchaete tenera, Cyllindrospermum muscicola* and others represent the major microorganisms which contribute to soil fertility (Abdel-Raouf N. *et al.,* 2012). The microalgae, a group of evolutionary old group of living organisms on the Earth, represent a large-scale .gene memory“ of the plant kingdom. The photoautotrophic microalgal cells are mostly interesting organisms as a source of adaptive plasticity to changes in environment. In the unicellular algal population communication network between cells could exist before occurency and maintain cell-cell connection leading to the multicellular phenotype such as displayed at charophytes, for example soil filamentous cyanobacterium *Klebsormidium flaccidum* (Hori *et al.,* 2014). The multicellularity phenomenon is based on the composition of cell wall (Domozych and Domozych, 2014). The evolution hypothesis, that the Charophytes shifted to adapt in terrestrial habitats by production and branching ability of rhizomes connecting with the SLs production was reported (Ruyter-Spira and Bouwmeester, 2012). Only few reports bring out some information about strigolactones production in macro or microalgae. Delaux *et al.* (2012) reported no possibility to find SLs in microalgae due to lack of some genes of the SLs biosynthetic pathway. In the study of Delaux *et al.* (2012) SLs production was suggested only in case of branched macroalgae, such as *Nitella* and *Chara* (freshwater species). The occurence of the key proteins of SLs pathways describing for vascular plants were analysed in representatives of chlorophyte green alga, Zygnematales, the Charales, the Bryophytes etc. (Delaux *et al.*, 2012). These are the compounds: D27, carotenoid cleavage dioxygenases (CCD) CCD7 and CCD8 (Alder *et al.*, 2012). However, the phylogenetic analysis of the CCD sequences present in the databases from algae did not show the presence of such homologues, instead, only CCD7 and CCD8 homologues have been identified (Ahrazem et al., 2016). The CCDs are enzymes that are responsible for the oxidative cleavage of carotenoids at specific double bond to generate apocarotenoides. These enzymes are found in animals, plants, photosynthetic bacteria, algae and cyanobacteria. C_13_ volatile apocarotenoids produced by marine macroalgae exhibit growth-regulating properties (Baldermann *et al.,* 2013). Cyanobacterial CCDs from genera such as *Synechocystis* and *Nostoc*, as well as *Anabaena*, have not been well studied yet. Heo *et al.* (2013) have doccumented abundancy of 3 CCDs (NSC1-3) in cyanobacteria *Nostoc* sp. PCC7120. The NSC3 cleaves β-apo-8´-carotenal, NSC2 moreover β-carotene, NSC1 cleaves different substrates of bicyclic and monocyclic carotenoids. This point outs, that the enzymatic cleavage reactions of carotenoides or apocarotenoides catalyzed by various CCDs (Walter and Strack, 2011) are the most important to determinate novel SL-related compounds in algae.

Only a few studies have been conducted with SLs or SL-related compounds in algae. The main objective of this study was investigation of the microalgal strains that could be suitable for biotechnological purposes and of the agricultural importance (application benefit for the crop) in pest management including suicidal germination of parasitic weeds.

## Materials and methods

### Plant material - screening

Set of microalgae sp. obtained from algal collections (CALLA, CAUP, Czech Republic): *Amorphonostoc* sp., *Anabaena variabilis*, *A. elipsosporum*, *Calothrix* sp. *Coleastrum* sp*., Cylindrospermum alatosporum* CCALA 998*, Eustigmatos* sp*., Fisheriella* sp.*, Haematococcus pluvialis, Chlamydomonas reinhardtii (2 strains), Chlorococcum* sp*., Chlorogloea* sp*., Chlorokybus atmophyticus, Klebsormidium flaccidum, Leptolyngbia* sp., *Microthamnion strictissimum*, *Nanochlorpsis* sp., *Nostoc calcicola, Nostoc muscorum, N. ellipsosporum, N. linckia f. piscinale, N. commune, N.* sp. *CM (symbiont with GUNNERA), Oscilatoria limosa*, *Phormidium* sp*., Scotiellopsis terrestris, Stichococcus bacillaris, Symploca thermalis, Synechocystis* sp.*, Tribonema vulgare, Trichormus variabilis, Trochydiscus* sp., *Vicheria sp.,* the symbiotic algae with fungi in lichens - *Trebouxia sp. Asterochloris* sp.*, Trentepohlia aurea,* and the algae obtained by free natural sampling: freshwater macroalga - *Cladophora* sp. and *Chara* sp. (2 isolates), the freshwaterweed *Elodea canadensis* (intermediate between macroalga and plant) and the marine macroalgae - *Codium* sp*., Gracilaria dura, G. Bornea, G.* sp*., Cystoseira* sp., *Ulva lactuca* were screened. The algal biomass was produced in cultivation tubes with volume from 500 to 2000 ml according to experiments. The algal biomass was separated by centrifugation (4500 rpm, 5 min) from the medium, freeze dried and stored at room temperature until testing. The natural samples of algae and the waterweed *E. canadensis* were dried at room temperature. Some of the isolates were tested immediately after collection.

In vitro axenic culture of *Trebouxia arboricola* (University of Gdańsk, Poland, strain XPAL450 isolated from *Xanthoria parietina*) and *Trebouxia erici* (strain UTEX911, USA = *Asterochloris erici* strain H1005, CAUP Czech Republic) were used for the optimalization of the germination bioassays. The strain UTEX911 was isolated from lichen *Cladonia cristatella* from soil (Massatchusetts USA 1958). The both strains were batch cultivated in Erlenmayer vessels (125 ml) on shaker (150 rpm) at illuminition 200 µE.m^-2^.s^-1^ (cool-white fluorescent lamps) at 23 ^o^C for 14 days. The cultures were cultivated without source of anorganic phosphorus for the last 3 - 4 days. The growth of the *Trebouxia* culture was enhanced by addition of 1% glucose (w/v) to the culture medium (Zachleder and Šetlík, 1982), pH 7.8.

### Preparation of crude extracts for screening

The 200 mg of dry biomass (freeze dried) or fresh algal samples were extracted by 3 ml of organic solvents: 70% (v/v) methanol, 100% acetone or 100% ethyl acetate, in mortar and pestle with quartz sand. The homogenate was allowed to stand for 20 min in closed tube and centrifugated at 4500 rpm for 5 min and at 4 ^o^C. The extraction was repeated twice, extract combined and stored at - 20 ^o^C until use.

### Germ tube branching bioassay on the AM fungus Gigaspora rosea

Arbuscular mycorrhizal fungus *Gigaspora rosea* T.H. (Nicholson and Schenck, 1979) was used in the experiment. Treatment solutions were prepared by dilution of algal extracts in a ratio of 1:75 (v/v) with sterile distilled water. The 50 mm plastic Petri dishes were placed on 50 mm nitrocellulose membranes Pragopor (Pragochema, Czech Republic) supported by 70 mm cellulose discs, each saturated with 1500 μl of the test solution. The prepared dishes were left open for one hour at room temperature. Subsequently, 10 - 13 spores of AM sponges were inoculated on each nitrocellulose membrane using an automatic pipette. Petri dishes were placed in a humid chamber and incubated for 8 - 21 days at 28 °C. For evaluation, both nitrocellulose membranes and cellulose discs were perfused with a solution of 5 % ink (v/v) and 5% vinegar (v/v) in water. Spinning spores were counted using a preparative microscope and the result is given as a percentage of germinated spores.

### Germination of seeds of parasitic plants - in vitro bioassay

*In vitro* bioassays were performed by methods described in Matusova *et al.* (2004). The seeds of *Phelipanche aegyptiaca* and *P. ramosa* were surface sterilized in 2% (v/v) solution of sodium hypochlorite containing 0.02% Tween-20 (v/v) for 5 min and rinsed several times with sterile distilled water. After surface sterilization the seeds of *Phelipanche* spp. were placed onto glass fiber filter paper discs (120 mm in diameter, approximately 50 seeds per disc) onto wetted filter paper in Petri dishes, sealed with parafilm and the seeds were preconditioned at 21^o^C for 12 days in the dark in the growth chamber. After preconditioning phase, discs with the seeds were transferred onto discs with testing solution (“sandwich”) in new Petri dishes. Preparation of treatment solution was based on a volume 500µl of concentrated crude extract. The solvent was evaporated under vacuum pump, disolved in 100µl of acetone and diluted 10 or 100 times by distilled water. The experimental aliquote of 40µl of the tested algal extracts were applied on each disc. For each bioassay, distilled water and 0.01 or 0.001 mg.l^-1^ GR24 were negative and positive controls, respectively. The bioassays were repeated twice or three times. The germination of seeds was evaluated using light stereomicroscope (Carl Zeiss Jena, Germany), percentage of germination was calculated as number of germinated seeds from the total number of seeds.

### Roots analysis of Arabidopsis thaliana - in vitro bioassay

The *Arabidopsis thaliana* seeds (wt Col 0) with 10% (v/v) sodium hypochloride and drop of Tween 20 for 5 min were sterilized. The seeds were 3-times washed with sterile distilled water and stratified for 3 days at 4 °C. 75 ml of MS medium (Murashige and Skoog, 1962) containing 1% agar (w/v), 1 g.l^-1^ sucrose, pH 5.8 was poured into Petri plate (diameter 9 cm). The 6 seeds per plate were placed on the cultivation medium from the top margin of 1.5 cm. The excess of water was evaporated and the Petri plates were vertically standed at an 45° angle at illuminition 200 µE.m^-2^.s^-1^ (cool-white fluorescent lamps) and temperature 22±1 ^o^C. The root systems were evaluated after 8 days by scanning of images (Scanner, 1200dpi, 24bit), and root Analyzer (NIS-ELEMENTS, ver.3.22, LIM Prague, Czech republic).

### Short-term pots experiments

In first experiment, buds outgrowth measurements were performed on rms-1 (*ccd8*) mutant of pea (*Pisum sativum* L.) cv. Térèse (Rameau *et al.,* 1997). 120 µl of crude extract of microalga *Trebouxia arboricola* (extraction of 5 ml of fresh pelleted biomass, representing 100 ml of fresh culture was extracted by 6 ml ethyl acetate) and lichen *Xanthoria parietina* (1 g dry biomass was extracted by 6 ml ethyl acetate) were prepared and the crude extracts were diluted ten times (T/10 and X/10). For the treatment solution there was taken 5 µl of each crude extract into 5 ml of physiological buffer (2% PEG + 50% ethanol + 0.4% DMSO + 0.1% acetone). In precultured pea plants (3 nodes), the first two buds of young seedlings and apex were cutt. The 10 µl of final treatment solution on the upper intact bud was applied. The physiological buffer and physiological buffer including 1 µmol.l^-1^ GR24 were used as negative and positive controls.

In second experiment, the seeds of pea cv. Terno were soaked by algal homogenate of *T. arboricola* (strain XPAL450). The algal homogenate in centrifuged tube prepared from pellet and 17 ml of distilled water was mixed with 20 pea seeds for 24h in dark at room temperature. The treated seeds were germinated, ten seeds per pot. The pea plants were cultivated in 7-liter containers containing commercial garden soil (Agroprofi Garden, Agrocs Ltd., Czech Republic) and vermiculite (1:1). In experiment with AMF, the 15 g dose spore inoculum of *Rhizophagus irregularis* (Symbiom Ltd., Czech Republic) was incorporated into substrate mixture. All plants were grown in greenhouse at 22±3 ^o^C with the 16h photoperiode. The plants before flowering (mean 8 - 9 produced nodes per plant) were evaluated by germination rate, by determination of length (cm), fresh and dry weight of shoots (g), respectively.

### Mass Spectrometry Imaging of carlactone and its derivatives

The cell cultures of the alga *T. arboricola* ± P-free were rinsed and concentrated by centrifugation. The 2μl of cell suspension was loaded onto a nylon membrane Nylon 66, 0.2 μm (Supelco, Bellefonte, PA). Nylon membrane was fixed to the glass slides (Prosolia, Indianopolis, IN) by the means of doubled-side tape. DESI imaging analysis was performed using an OrbiTrap Elite (Thermo Fischer Scientific, Bremen, Germany) with a DESI-2D ion source (Prosolia, Indianopolis, IN). Imaging experiments were performed by continuous scanning of the surface. Spraying liquid (acetonitrile / 0.1% acetic acid mixture, v/v) at flow rate 3 μl.min^-1^, scanning velocity 65 μm.s^-1^ and an 65° spray impact angle were used. Data were acquired in the mass range m/z 50 - 800. Typical time of the analysis was less then 120 min. The obtained data were processed by the means of the BioMap software and two-dimensional ion images were created. Parameters of the MS analysis were optimised to following values: nebulizer pressure (N_2_): 7 bar, capillary heating: 300 °C, spray voltage: 5 kV, lens voltage: 60 V, ion injection time: 400 ms. Two microscans were carried out for each pixel. DESI-MSI analysis was performed in positive ion mode for carlactone (*m/z* 303.195), methyl carlactonoate (*m/z* 347.184), carlactonoic acid (*m/z* 333.169), 19-hydroxy carlactone (*m/z* 319.191) and 19-oxo carlactone (*m/z* 317.173).

### DART ambient technique

DART-Standardized Voltage and Pressure Adjustable (SVPA) ion source with tweezer holder module (IonSense, Saugus, MA) was coupled to Orbitrap Elite mass spectrometer (Thermo Fischer Scientific, Bremen, Germany) through the interface evacuated by the diaphragm pump. The DART ion source was operated in the positive ion mode with helium ionizing gas at the pressure 0.65 MPa. The beam was heated in the temperature range 300 °C to 400 °C, the grid electrode voltage was in the range of 300 - 350 V. The parameters of the mass spectrometer were following: capillary voltage 50 V, tube lens voltage 100 V, skimmer voltage 18 V and capillary temperature in the range of 300 °C – 350 °C. The acquisition rate was set to 2 spectra.s^-1^ with mass resolving power of 120,000 FWHM. All DART mass spectra were acquired over a mass range of *m/z* 50-400. Xcalibur software (Thermo Fischer Scientific, Germany) with DART web-based module was used for the instrument operation, data acquisition and processing.

### RNA extraction and marker gene expression - Ppccd8 mutant of Physcomitrella patens

The *Ppccd8* mutant (Proust *et al.,* 2011) has been used for the experiment. Transcript levels of SL response markers - PpCCD7 and Pp3c2_34130v3.1 homologous to KAR-UP F-BOX1 (KUF1) a SL response marker from *Arabidopsis* (Nelson et al. 2011, Lopez-Obando and Bonhomme, unpublished data) 6 or 16h after treatment based on published method (Lopez-Obando *et al.,* 2016) were obtained. Total RNA was extracted using QIAGEN RNeasy mini-kit with the column DNAse treatment. Absence of DNA contamination was checked by PCR. CDNAs was prepared from 2 µg of each sample RT-qPCR for PpCCD7, Pp3c2_34130v3.1, *PpACT3* and *PpAPT* genes. For gene expression analysis, genes were normalized against the mean of *PpACT3* and *PpAPT* genes (Lopez-Obando *et al.,* 2016). Preparation of dry ethyl acetate extracts from: microalgae *T. arboricola* grown in complete medium *(Trebouxia)* and in P-free medium for last 3 days of culturing (*Trebouxia-*P): 5 ml of fresh pelleted biomass, representing 100 ml of fresh culture, was extracted by 6 ml ethyl acetate; lichen *X. parientina* (*Xanthoria*): 1 g dry biomass was extracted by 6 ml ethyl acetate and 200 ml of exhausted culture medium of *T.arboricola* (Medium *Trebouxia*) was concentrated on silicon column C18 and eluated by 500 µl ethyl acetate. The crude extracts were stored at -20 ^o^C. Preparation of dry samples and treatments solutions: 1000 µl ethyl acetate of each extract was evaporated in vacuum, dry samples were resuspended in 120 µl of acetone and diluted 1:9 with acetone. Bioactivity of 25 µl tested extracts of *Trebouxia, Trebouxia*-P*, Xanthoria,* Medium *Trebouxia* and 10 times diluted the samples (T/10, X/10, MT/10 and T-P/10) on *Ppccd8* mutant grown for 6 or 16 hours from spores on 25 ml minimal PP-NO_3_ medium (Hoffman et al. 2014, Ashton, 1979) were applied. The 25 µl of acetone (negative control), 1 µmol.l^-1^ GR24 (positive control) were used.

### Statistical analysis

Results were analysed by analysis of variance Anova test *p* < 0.05, Statistica software Statistica ver.8.0 (StatSoft Inc. USA) followed by Tukey´s and Kruskal-Wallis significance tests at the 5% level.

### Results and discussion

Two antagonistic roles of SLs in the rhizosphere are known: 1) they facilitate the formation of symbioses with arbuscular mycorrhizal fungi, rhizoid elongation and branching, mainly under P-limitation; 2) they are signals for the parasitic weeds, such as the *Striga, Phelipanche* and *Orobanche* species, indicating the presence of a host species, resulting in devastating losses in some agricultural systems. Our study focused on cultureable strains of algae that potentially could help to eliminate parasitic weed seeds from the soil. First, we tested physiological effects of algae on germination rate of the parasitic seeds, on roots growth and AMF spores and mycelium development in different bioassays. Then we focused on detection and determination of SLs biosynthetic genes and SLs in algae. Finally, were tested the selected algal strain in pots experiment.

### Screening of algae and plants for bioactivity - germination of seeds of parasitic plants

The list of tested algae and plants represents a random selection in developmental line and it includes representatives of various families, especially green coccals, filamentous algae and cyanobacteria, representatives of marine and freshwater branched macroalgae, which are expected to synthesize SLs. Crude 70% (v/v) metanol, ethyl acetate or acetone extracts of algae (freeze-dried, dried or fresh, **Table 1**) were prepared. The results of germination bioactivities suggested an independence from position of developmental line (cyanobacteria, microalga, macroalga, marine or freshwaterweed) and filamentous or branched types. Our test using germination bioassay and ethyl acetate extraction have shown on the same bioactivity to SLs in filamentous freshwater Charales (*Chara* sp.) and Ulvophyceae (marine *Ulva lactuca*, freshwater *Cladophora* sp.) macroalgae as was reported by Delaux *et al.* (2012). In addition the stimulation bioactivity of other marine branched macroalga *Cystoseira* sp., *Gracilaria* sp., *Codium* sp., freshwaterweed *Elodea canadensis* was found.

**Table 1.** The germination of *Phelipanche aegyptiaca* seeds, induced by crude ethyl acetate extracts of selected microalgae macroalgae and freshwater plant (screening). In bold – filamentous or branched type alga. Stimulating germination bioactivity of tested alga extract > H_2_O (negative control); 0.001 mg.l^-1^ GR24 (positive control). Germination seeds in %.

| **Blue-green alga** | no activity | stimulating activity | **Macroalga** | stimulating activity |
| --- | --- | --- | --- | --- |
| Cyanophyceae | *Amorphonostoc* sp.<br>*Anabaena variabilis*<br>*A.elipsosporum*<br>*Calothrix* sp.<br>*Leptolyngbia* sp.<br>*Fischeriella* sp. | *Cylindrospermum alatosporum*<br>(soil)<br>(17% ethyl acetate) | Ulvophyceae | *Cladophora* sp.<br>(45% ethyl acetate)<br>*Ulva lactuca*<br>(32% ethyl acetate) |
| Nostocaceae | *Nostoc muscorum*<br>*N. ellipsosporum*<br>*N.linckia f. piscinale*<br>*N. calcicola*<br>*N.commune*<br>*Trichormus variabilis* | *N. commune* (symbiont with GUNNERA)<br>(2.5% ethyl acetate) | Fucaceae | *Cystoseira* sp.<br>(28% ethyl acetate) |
| Oscillatoriaceae | *Oscillatoria limosa* |  | Gracilariaceae | *Gracilaria dura*<br>(49% ethyl acetate)<br>*G .bornea*<br>(34% ethyl acetate) |
| Phormidiaceae | *Phormidium cruentum*<br>*Symploca thermalis* |  | Charophyceae | *Chara* sp.<br>(29% ethyl acetate) |
| Chroococcales | *Synechocystis* sp. |  | Bryopsidales | *Codium* sp.<br>(13% ethyl acetate) |
| **Microalga** |  |  | Hydrocharitaceae | *Elodea* sp<br>(35% ethyl acetate) |
| Trebouxiaceae | *Microthamnion strictissimum*<br>*Stichococcus bacillaris* | *Asterochloris* sp., *Trebouxia arboricola*, *T. erici* (symbiont)<br>(33% ethyl acetate) | **Lichen** | *Xanthoria parietina*<br>(56% acetone),<br>(13% ethyl acetate) |
| Trentepohliaceae |  | *Trentepohlia aurea* (32% ethyl acetate) |  |  |
| Chlorophyceae |  | *Chlamydomonas reinhardtii*<br>(26% ethyl acetate) |  |  |
| Chlorococaceae | *Chlorococcum* sp. |  |  |  |
| Eustigmatophyceae | *Trochodiscus* sp.<br>*Vicheria* sp.<br>*Nanochloropsis* sp. | *Eustigmato* sp. (symbiont with *Chara*, water)<br>(37% ethyl acetate) |  |  |
| Haematophyceae | *Haematococcus pluvialis* |  |  |  |
| Chlorophyta | *Chlorogloea* sp. |  |  |  |
| Klebsormidiaceae | *Chlorokybus atmophyticus*<br>*Klebsormidium flaccidum* |  |  |  |
| Tribonemaceae | *Tribonema vulgare* |  |  |  |
| Scenedesmaceae | *Coleastrum* sp.<br>*Scotiellopsis terrestris* |  |  |  |

The most important is the extraction procedure and the type of extraction agent. In sample of the *Cladophora* sp., germination inducing compounds were not extracted in 70% methanol, in comparison to ethyl acetate extract (**Table 2A**). Methanol as extraction agent was excluded for germination bioassays. Based on preliminary *P. aegyptiaca* germination bioactivity screening of macroalgae, lichens and water plant extracts, we futher tested crude extracts obtained by ethyl acetate extraction, concentrated by evaporation of solvent, dissolved in small amount of acetone and dilluted with water (1:9 v/v) (**Table 2B**).

**Table 2A.**
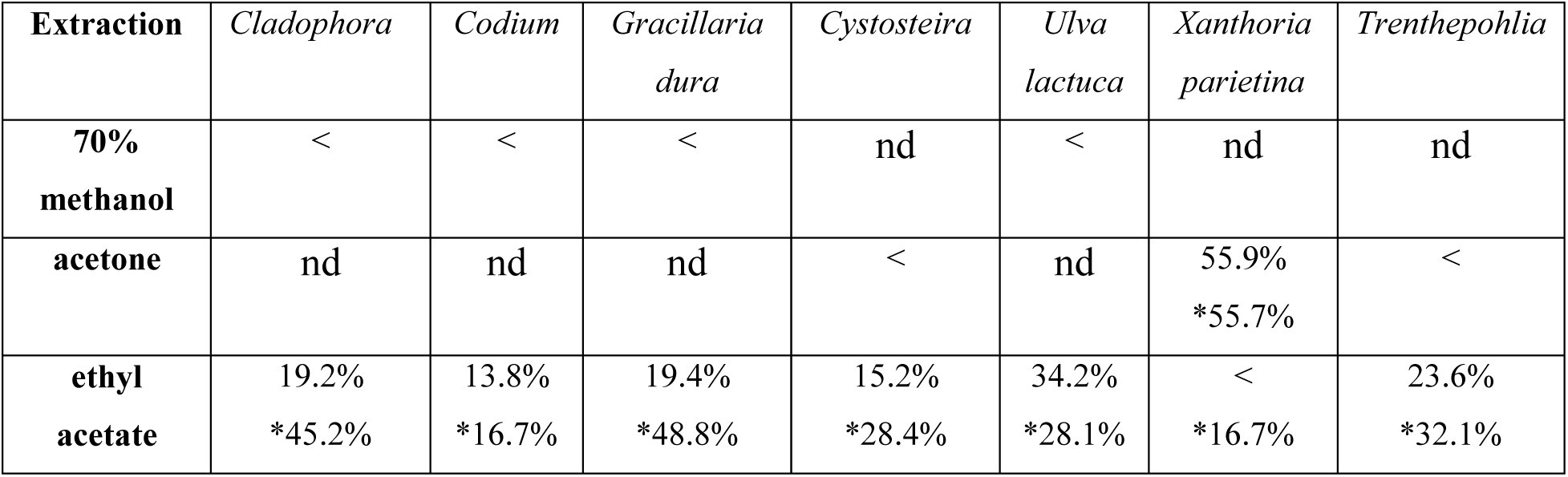
Germination of *Phelipanche aegyptiaca* seeds induced by crude extracts of algae. Algae were extracted in 70% methanol, acetone and ethyl acetate. Germination (%) of crude extracts and extracts 10x diluted (*), H_2_O was used as a negative control (12.5±4.2%) and 0.001 mg.l^-1^ GR24 as a positive control (75.0±4.0%); nd - not determined. Germination seeds in %.

**Table 2B.**
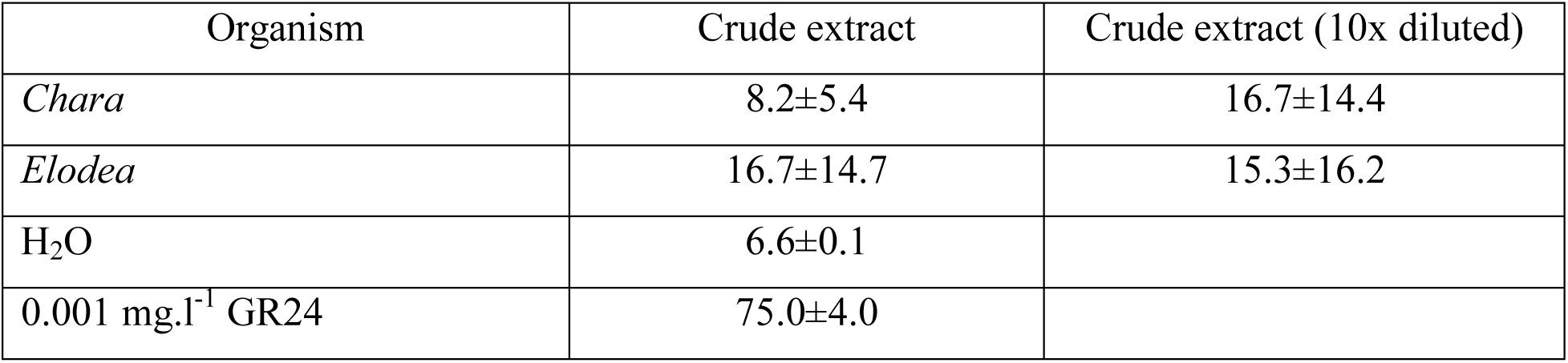
Induction of germination of *Phelipanche aegyptiaca* seeds by macroalga *Chara* sp. (33gfw/45ml) and freshwaterweed *Elodea canadensis* (47gfw/70ml) crude extracts (ethyl acetate). Germination seeds in %, mean±SD.

Futher we found that two soil cyanobacteria filamentous strains (soil and symbiotic) and four microalgae (symbiont and one with flagella) stimulated the germination of the *P. aegyptiaca*. They were *Cylindrospermum alatosporum* (CCALA 998) and *Nostoc commune* (symbiont with roots of Gunnera). The soil representatives of cyanobacteria form a spherical cell-free colony of cells, rich in carotenoids and able to N-fixation in heterocyst without the presence of oxygen. The lichenized algae: the carotenoid-rich red microalga *Trenthepohlia aurea* is lichen growing on tree and rich on specific pigments, the most common photobiont genera *Asterochloris* and *Trebouxia* (green microalge cultures grown without fungi) as symbiont of many lichens. The other two selected strains *Eustigmatos* sp. water isolates from *Chara* sp. and flagellate green freshwater alga *Chlamydomonas* sp. stayed independent. In the study Delaux *et al.* (2012) cyanobacteria was not tested, but Heo *et al.* (2013) reported unique cleavage activity of NSC3, a CCD of *Nostoc* sp. strain PCC 7120.

The information about presence of genes associated with the biosynthetic pathway of strigolactones in microalgae is still missing. Delaux *et a.,* (2012) reported presence of putative homologous proteins of receptor D14 in *Klebsormidium* sp., CCD8 and receptor D14 in *Chlorokybus* sp. and CCD8 in *Trebouxiophyceae*. We tested the hypothesis that SLs could be also produced in microalgae. Unlike *Trebouxia* sp. and *Asterochloris* sp. at both common soil charophytic algae *Chlorokybus atmophyticus* and *Klebsormidium flaccidum* germination bioactivities for *P. aegyptiaca* were not observed. However, we can not rule out the inhibition of SLs biosynthesis in green algae (Delaux *et al.*, 2012) or specific germination induction of other parasitic weed seeds. More detailed study using other parasitic weed seeds and analytical approach may help to answer this question. In addition, *K. flaccidum* synthesizes several plant hormones and gained many genes typical for land plants (Hori *et al.,* 2014).

The branching function of SLs is accounted as original function in plants, in moss *Physcomitrella patens* (regulation of protonema branching), in Charales and in fungi. In *P. patens* SLs act as reminiscent sensing molecules used by bacteria to communicate one with eachother (Proust et al. 2011). In unbranched filamentous microalgae and cynobacteria, there is a mutual communication of individual cells as was reported also for green algae such as multicellular *Volvox globator* or the charophyte (Domozych and Domozych, 2014). A specific group, where two different kinds of organisms - algae and fungi communicate, are lichens. In our experiment we tested cyanobacteria, lichenised algae and lichen for bioactivity similar to SLs. **Fig. 1** shows the results of induction of germination of *P. aegyptiaca* seeds in the bioassay with crude extracts of *Trebouxiophyceae* algae (lichens photobiont from artificial culture). All the algal treatments increased the germination activity of the seeds. GR24 (100µmol.l^-1^) treatments added to the alga extract restored the germination bioactivity similar to the GR24 alone treatment, this means that the algal extract didn´t contained substances that inhibit GR24 effect. It seems, that lichen and linechised algae contain SL-related compounds. We need to further optimise growing conditions, and/or an extraction procedure.

**Fig. 1.**
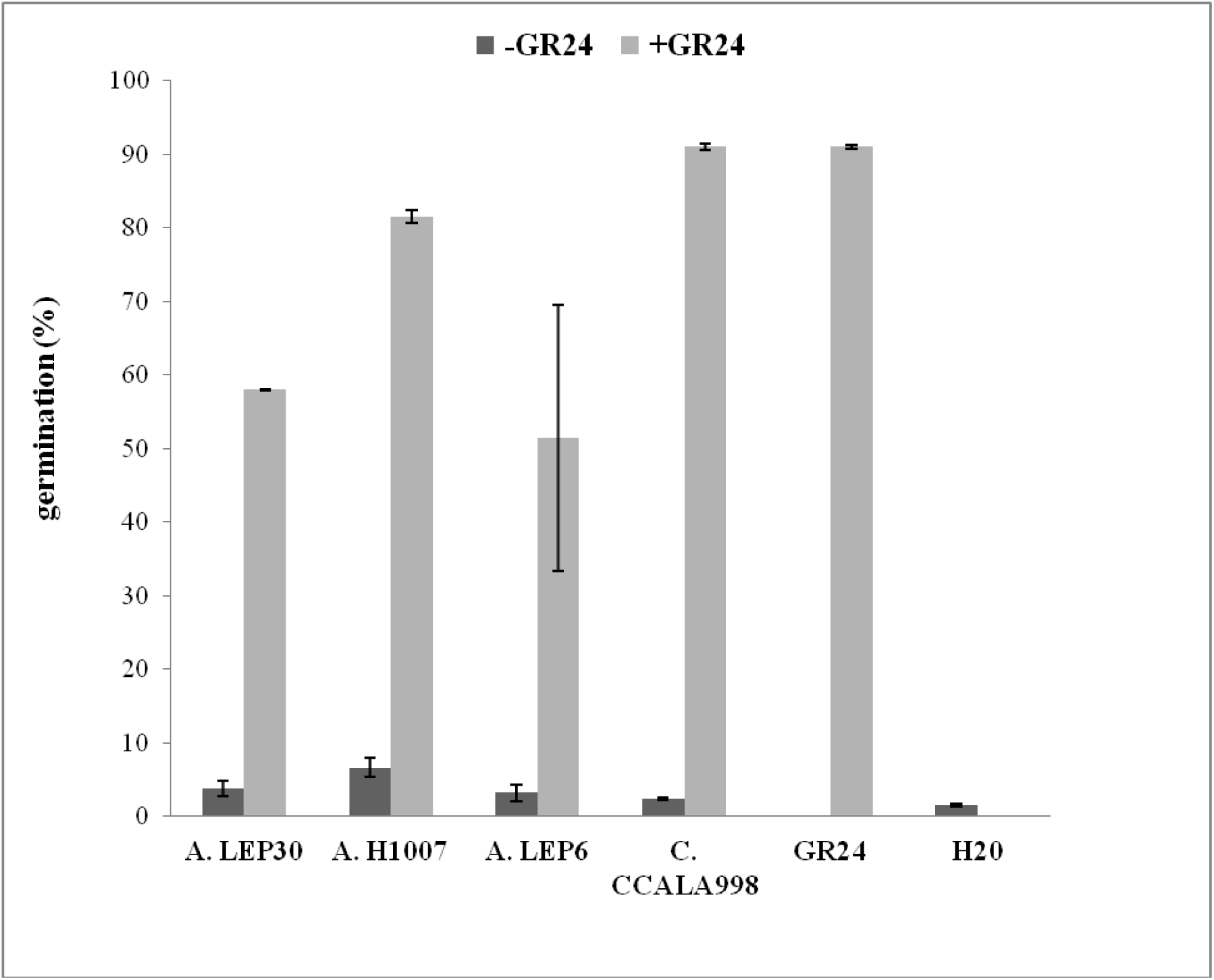
Germination of *Phelipanche aegyptiaca* seeds. Ethyl acetate crude extracts of green algae - *Asterochloris* sp. strain LEP30, *A. excentrica*, strain H1007, *A*. sp. strain LEP6 and blue-green alga - *Cylindrospermum alatosporum* strain CCALA998 were applied on preconditioned seeds. Extract treatments 7^th^ days and evaluation of germination rate in %. After 7 days GR24 (100 µmol.l^-1^) treatment to the tested algae restored the germination bioactivity. Values are mean ± SD of 2 replicates.

### Trebouxia cultures – optimalization of in vitro bioassays

Based on the screening and the data of biostimulation activity of common lichen *Xanthoria parietina* and their photobionts *Trebouxiophyceae* algae (*Trebouxia* sp. syn. *Asterochloris* syn. *A. excentrica* UTEX1714 (USA); *A. erici* UTEX911 from lichen *Cladonia cristatella* (USA), *A.* sp. LEP30 (CZ), *T. arboricola* (PL) on the seeds of P*. aegyptiaca* we supposed, that the lichenized alga produce SL-related compounds or compounds with SL-like bioactivity. **Table 3** show data of germination rate (%) of seeds of two broomrapes *P. aegyptiaca* and *P. ramosa* in a presence of crude ethyl acetate extract of *Trebouxia* sp. (symbiont, photobiont in lichen) which again suggested presence of SL-related compounds in the alga. Exhausted medium probably also contains SL-related compounds. The cells exudates were concentrated on C18 column, eluated with acetone or ethyl acetate similar to Kohlen *et al.* (2011). Stress conditions in plants in view of SLs production exhibited increase of SLs production (Yoneyama *et al.,* 2012). Based on this knowledge we tested P-free culture of the *Trebouxia arboricola* (see Material and methods) on the efficiency in germination bioassay. Ethyl acetate optimized extract from *T.arboricola* P-free dry weight biomas led to increase of germination rate in *P.aegyptiaca*. P-free medium cultivation and used extract i.e. the actual concentration of SLs in the applied sample, have the high significancy on the germination rate. Delaux *et al.* (2012) used three times extraction of fresh tissue (10 - 80g) of algae by acetone and the extract was dried and dissolved in ethyl acetate. Yoneyama *et al.* (2012) used the extraction of roots by acetone. Furthermore at both studies, the extracts were washed with 0.2M KH_2_PO_4_, dried over anhydrous MgSO_4_ and concentrated in vacuo and stored at -20 ^o^C for AMF bioassay and determination of SLs in extracts. In our procedure the first extraction was carried out with ethyl acetate, then the extract was evaporated to dryness and stored at -20 °C. Just before bioassay, the samples were dissolved in acetone and dilluted with sterile distilled water. SLs are acting at picomolar to nanomolar concentrations (Akiyama and Hayashi, 2006). Therefore the optimization of extraction procedure was tested in the bioassay with *P. aegyptiaca seeds.* Different dilutions of crude extracts were prepared. Starting from 5 ml of alga pelet, 3 ml of crude ethyl acetate extract was applied: in 10x dillution with acetone (extract 1), 10x diluted with water (extract 2), 2.5x concentrated (extract 3) and extract 3 was 10x diluted with water (extract 4). Extract 1 induced 4.70 ± 5.74% germination of *P. aegyptiaca* seeds, extract 2 induced 4.73 ± 4.27% germination, extract 3 induced 7.83 ± 2.23% germination and extract 4 13.03 ± 8.01% germination. The controls of germination were 0.0 ± 0.0% (H_2_O) and 77.0 ± 6.79% (GR24, 0.01 mg.l^-1^). The observed increase in germination of seeds treated by the *Trebouxia arboricola* suggested in symbiotic microalgae SLs or SLs-like activity compound(s) production.

**Table 3.** Stimulation activity in the selected group of *Trebouxiophyceae* algae for germination bioassays with *Phelipanche* spp.. Optimization of preparation extract of alga *Trebouxia arboricola* (etylacetate:5mlfw/6ml for both *P*. spp.) and extract of cultivation medium (etylacetate:1000mgdw/8ml). Optimization of cultivation by P-free medium in stationary phase of growth - extract of alga *T. arboricola* (etylacetate:360mgdw/8ml). Extract of lichen *Xanthoria parietina* (etylacetate:1000mgdw/9ml). Germination seeds in %, mean±SD.

| Testing agent | *P. aegyptiaca* | 10x dilution | Testing agent | *P. ramosa* | dilution |
| --- | --- | --- | --- | --- | --- |
| *Trebouxia* | 7.2±0.4 | 42.5±9.2 | *Trebouxia* | 18.6±1.4 | 9.0±3.6 |
| medium *Trebouxia* | 4.2±5.9 | 11.5±3.5 | H<sub>2</sub>O | 3.6±3.4 | - |
| *Trebouxia-P* | 19.4±0.6 | 11.8±0.3 | 0.01 mg.l<sup>-1</sup>GR24 | 96.3±0.1 | - |
| *Xanthoria* | 20.1±3.4 | 10.0±2.5 |  |  |  |
| H<sub>2</sub>O | 6.0±0.6 | - | - |  |  |
| 0.001 mg.l<sup>-1</sup>GR24 | 71.3±3.8 | - | - |  |  |

### Roots analysis of Arabidopsis thaliana - in vitro bioassay

The plants were grown in Murashige and Skoog medium (1962) in presence of crude extracts. We measured total length of roots, length of primary root and length of lateral roots of 8-days old plantlets of *Arabidopsis thaliana* after application of tested substances. Roots analysis (**Table 4**) documented the statistical significant biostimulation of *T. arboricola* crude extract in roots. The other crude extracts of dry lichen *Xanthoria parietina*, exhausted *T. arboricola* medium concentrated on C18 column and eluated by acetone, freshwater macroalga *Chara* sp. or freshwaterweed *Elodea canadensis* did not revealed any changes in measured parameters of roots. This bioassay also suggest presence of compounds with SL-like activity in the alga *T. arboricola*. Strigolactones, which are produced mainly by plant roots, affect plant root development and architecture via the control of cell division observed in wt *A. thaliana* (Koltai, 2011). Application of GR24 (10^-6^M) increases root hair elongation in wt *A. thaliana* (Ruyter-Spira *et al.,* 2011), but not at higher concentration of GR24 in tomato (Koltai, 2011). Results of our study suggested the increase in length of primary and lateral roots and repression lateral adventitious root formation (total number of roots) in wt *A.thaliana* after application of extracts from T. *arboricola* pellet. The physiological effects correspond to the effect of GR24 application (Sun *et al.,* 2016).

**Table 4.** Evaluation roots of 8-days old seedlings of *Arabidopsis thaliana* (wt, Col-0) growing in 1/2 MS medium (half strength of Murashige and Skoog medium) with crude extracts of *Trebouxia arboricola* pellet, exhausted cultivation medium of *T. arboricola*, biomas of freshwater macroalga *Chara* sp., biomas of freshwater higher water plant *Elodea* sp. and lichen *Xanthoria parietina.* Evaluated parameters by image analysis: TL – total length of roots; T – total number of roots; PL – length of primary root; LL – length of lateral roots. Mean values ±SD marked with distinct letters differ significantly one another according to Tukey´s test (p < 0.05).

| Sample | TL[mm] | T | PL[mm] | LL[mm] |
| --- | --- | --- | --- | --- |
| CONTROL | 38.89±7.8<sup>abc</sup> | 4.6±1.7 | 27.7±2.7<sup>c</sup> | 11.2±6.3 |
| *Trebouxia* | 45.68±9.8<sup>ab</sup> | 4.1±0.9 | 32.6±3.2<sup>ab</sup> | 13.1±9.2 |
| *Xanthoria* | 36.69±9.0<sup>bc</sup> | 4.3±1.9 | 22.8±11.0<sup>cd</sup> | 13.9±6.8 |
| medium *Trebouxia* | 32.77±6.8<sup>c</sup> | 4.6±1.4 | 22.7±4.3<sup>d</sup> | 10.0±5.3 |
| *Chara* | 33.72±4.3<sup>c</sup> | 3.5±1.6 | 25.0±7.9<sup>cd</sup> | 8.7±8.8 |
| *Elodea* | 33.49±11.7<sup>c</sup> | 4.1±2.5 | 24.7±5.9<sup>cd</sup> | 8.8±7.8 |

### Germ tube branching bioassay on the AM fungus Gigaspora rosea, stimulation growth of fungi by Trebouxia arboricola

When AMF responds positively to the presence of algae extract, it is possible to predict the presence of SLs in the algae. The treatment of ethyl acetate crude extract of Charales alga *Nitella mucornata* and cyanobacteria *Cylindrospermum alatosporum* increased AMF spore germination (**Fig. 2A**). The treatment of acetone crude extract of lichens led to changes in the growth of mycelium of AMF *G. rosea* observed as increasing production of large mycelium (**Fig. 2B**). All of natural strigolactones are active as branching factor also in AMF *G. rosea* (Akiyama and Hayashi, 2006). There is further evidence that some algae and lichens (fungi or algae) can produce SLs or SLs-related compounds. As reported earlier, SLs stimulation effects on fungal hyphe growth in AMF (Besserer *et al.,* 2006, 2008) would probably exist in lichen symbionts. In lichens, SLs have been suggested as potential candidates for the photobiont-derived factor which induces hyphal branching of the mycobiont after the initial contact of alga and fungus (Harris, 2008). In lichens the fungi may be selecting very specific algal genotypes, while the algae are tolerant of many fungal partner (Piercey-Normore and DePriest, 2001). The hypothesis about existence of communication connecting with SL-related compounds between phycobiont (alga) and fungi needs to be further tested. Our preliminary data of the increased biomass of the culture beneficial fungi *Lecanicillium muscarium* in a presence of exhausted medium of *T. arboricola* supported the hypothesis about SLs production by the alga in the medium.

**Fig. 2.**
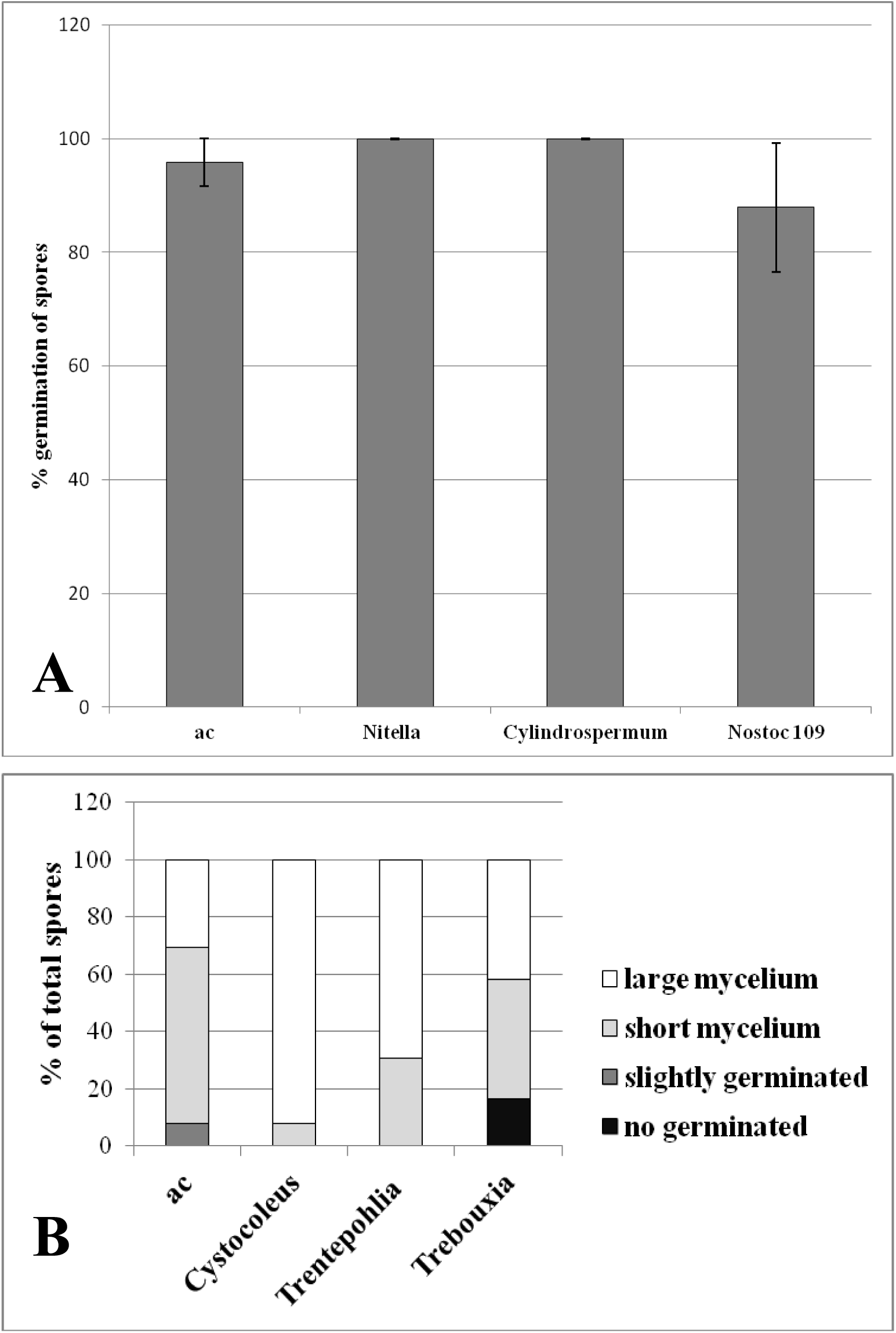
Germination of spores and growth of mycelium of AMF *Gigaspora margerita* effected by **A)** aceton extract of Charales: *Nitella mucornata* and blue-green algae: *Cylindrospermum alatosporum* (dw 0.23g); *Nostoc* sp. strain 109 (dw 0.22 g) and by **B)** aceton extract of lichens natural dry samples: *Cystocoleus ebenus* (dw 0.25 g), *Trebouxia* (dw 1.0 g) and free-living alga *Trenthepohlia* sp. (dw 0.25 g). Values are mean ± SD of 6 replicates.

### Expression of SL-related markers - Ppccd8 mutant of Physcomitrella patens

The *Physcomitrella patens* (moss) knockout mutant *Ppccd8* (Proust *et al.*, 2011) was used for our study as the response to SL (Proust *et al.*, 2011, Hoffmann *et al.*, 2014). On the **Supplementary Fig. S1** are shown the transcript levels in *Ppccd8* mutant of SL response marker genes PpCCD7 and Pp3c2_34130v3.1, respectively 16 and 6h after application of acetone ac (control), 1 µmol.l^-1^ GR24, alga *Trebouxia arboricola,* lichen *Xanthoria parientina*, exhausted culture medium of the alga *T. arboricola.,* T/10 and X/10. The results suggested effects of the algae treatment on the expression level of the marker genes. Although the effect of GR24 on PpCCD7 transcript levels are significantly dampened by *Trebouxia arboricola* diluted extracts, as it is expected from SL compound (Hoffmann *et al.*, 2014). Diluted *Trebouxia*, *Xanthoria* and diluted *Trebouxia-P* extracts all led to an increase of the Pp3c2_34130v3.1 gene transcript levels, as GR24. These results further indicated the presence of SLs-related compounds in microalga *Trebouxia arboricola* and lichen *Xanthoria parietina*.

Here, we present data indicating that symbiotic *Trebouxiophyceae* also produce SLs or SLs-related compounds, but we were unable to amplify CCD8 sequences from the genomic DNA for any screened algae due to limited genomic and transcriptomic data. Whital across screened 37 species of cyanobacteria for novel CCD genes used BLASTP, 5 CCD genes including *ccd8*-homologous gene in freshwater unicellular N_2_-fixing cyanobacteria *Cyanothece* sp.ATCC 51142 and more than 3 CCD genes including *ccd7*-homologous genes in filamentous cyanobacteria *Anabena* and *Nostoc* were identified (Cui *et al.* 2012). It is known, that both CCD7 and CCD8 were envolved due to the duplication of CCD1 genes in plant families, CCD genes should be functionally divergent from each other, the CCD7/8 genes had greatest distance between the mosses (*P.patens*) and other angiosperm species (Priya *et al.,* 2014). The last fact could be the reason, why it was not possible to find suitable nucleotide sequences to find CCD8 in microalga *Trebouxia*. Cui *et al.* (2012) summarized hypothesis about *ccd7*-homologous genes origin in cyanobacteria, while *ccd8*-homologous *genes* were absent because of gene loss. Another possible hypothesis is the fact that microalgae, especially soil microalga and cyanobacteria, are considered as pioneers in settling the Earth’s surface after, for example, a fire. This leads us to believe that these algae would still have unknown receptors, perhaps more similar to receptors for karrikins (Flematti *et al.*, 2009).

### Short-term pots experiments

In first experiment the effects of crude extract of microalga *Trebouxia arboricola* and lichen *Xanthoria parientina* and ten times diluted samples (T/10 and X/10) on the outgrowth of the lateral bud after cutting the main stem were examined. Lateral growth restoration was determined by measuring of the length of the outgrowing lateral shoots in comparison to controls – with and without treatment of 1 μM GR24 (**Fig. 3**). The inhibition of branching in rms-1 branched phenotype was expected after applications of *Trebouxia* or *Xanthoria*, respectively. The difference was very significant between control and GR24. Limit of significance between control and *Trebouxia* and between control and T/10, since the *p* value is superior to 0.05 but not far from it (*). The most effective remains inhibition by synthetic stimulant GR24, functioned in the experiment as a auxine analog (full inhibiton of outgrowth of lateral buds after cancelation of apical dominance). In plants, SLs production exhibited by declination of branching activity of shoots, ie inhibition of outgrowth of axillary buds (Umehara *et al.,* 2008, Kohlen *et al.,* 2011). This result assumes in the alga *Trebouxia* the existence of substances that have the same phenotypic response as the GR24 application. *Mass Spectrometry Imaging of carlactone and its derivates and fragments identification by using DART-HRMS*

**Fig. 3.**
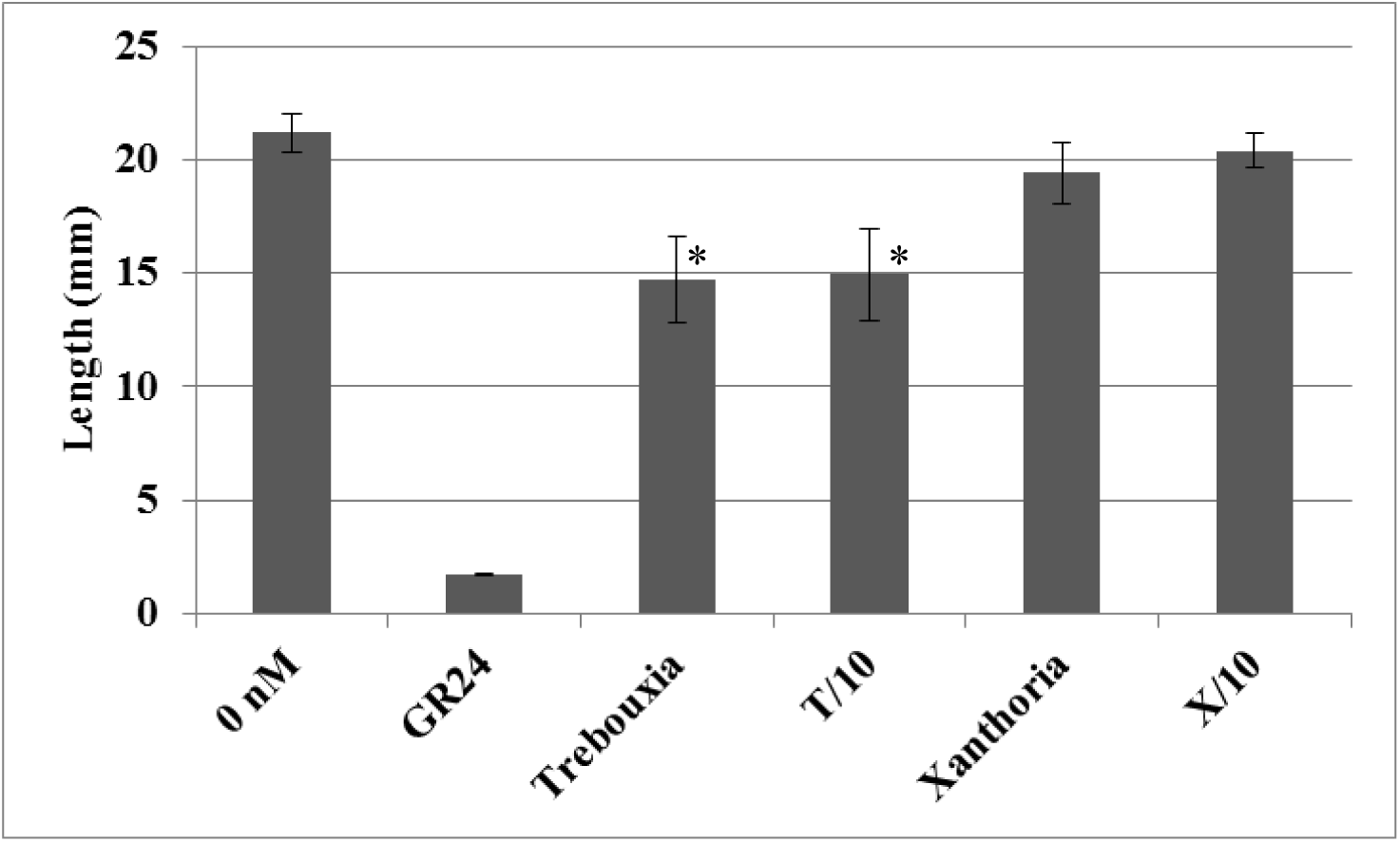
Effects of crude extract of microalga and lichen on the outgrowth of the lateral bud after cutting. Lateral growth restoration was determined by measuring as the length of the outgrowing lateral shoots. Crude extracts of *Trebouxia arboricola* (Trebouxia), 10x diluted extract of *T. arboricola* (T/10), *Xanthoria parientina* (Xanthoria), 10x diluted extract of *X. parientina* (X/10), 1μM GR24 as a positive control and physiological buffer (0 nM - containing 2% PEG, 50% ethanol,0.4% DMSO, 0.1% acetone) were used. Values are mean ± SD of 20 replicates. (*) indicate statistically significant differences from control (0 nM) according to Kruskal-Wallis test (P < 0.05, for *Trebouxia* P=0.019, for T/10 P=0.02).

Carlactone is being a key molecule in SLs biosynthesis (Seto *et al.*, 2014). DESI-MSI is a modern analytical technique that enables to measure real samples at ambient conditions, including highly troublesome analytes that are otherwise not easy to identify due to chemical changes during necessary sample preparation or extraction steps. To our knowledge, we present for the first time identification of carlactone and carlactone derivatives by this method. Our results present determination of β-apo-13-carotenone: carlactone, methyl carlactonoate, carlactonoic acid, 19-hydroxy carlactone and 19-oxo carlactone in green unicellular coccal microalga *Trebouxia* (free-cultivate cells). We found β-apo-13 carotenone, a second alternative to carlactone synthesis. 9-cis-β-apo-10'carotenal did not find, at least in a measurable area. It is possible to use it more for synthesis of carlactone and beta-apo-13 carotenone is more measurable because its conversion to carlactone is slower. CCD8 (EC 1.13.11.70) catalyzes conversion all-*trans*-β-apo-10´-carotenal into β-apo-13-carotenone and this reaction is slower than that with *cis* isomer (Alder *et al.*, 2012). DESI-MSI images (see **Fig. 4A,B**) show the localisation of the above-given analytes in the cells of *Trebouxia arboricola* from the harvested culture grown on full culture medium (A); and in P-free medium (B). The relative ion intensity values corresponding the colour coding can be found in the bar on the right side. It is shown that short term P-free growth of *Trebouxia* culture led to increase of carlactones content in the cells. This result agrees with the previous study of Yoneyama *et al.* (2012). The result supports our hypothesis that there is CCD8 homologous gene. If the primary function of SLs is regulation of branching (Ruyter-Spira and Bouwmeester, 2012, Brewer *et al.,* 2013) that the CCD8 gene could assist in the alga in a symbiosis with fungi similar to animal-cyanobacterial symbiont *Cyanothece* sp. PCC7425 (Cui *et al.*, 2012). They reported that CCD8 enzymes are present in all eukaryotes including algae, while absent in all cyanobacteria except the symbiont (obtain of the gene by horizontal gene transfer or under selection during evolution). The identity of individual carlactone derivatives was validates both by the means of accurate mass and via other method, DART-HRMS. This method enables validation via fragmentation, the results are given in **Supplementary Table S1**, data were obtained fom extract. By the means of the MassFrontier program, probable fragments of individual carlactone analytes were calculated according to the fragmentation rules were obtained. Occurrence of individual fragments was monitored in real samples. According to the obtained results, relatively good similarity was achieved between the theoretically calculated fragments and the determined ones. Values of the diference between measured and theoretical accurate mass varied in the range of 0.03 ppm –1.82 ppm. For the analytes that had already been studied in other matrices (carlactone, carlactonoic acid, methyl carlactonoate), comparison of fragmentation results was performed and the experimental data were found to be in a good accordance with previously published data (Seto *et al.*, 2014, Abe *et al.*, 2014, Brewer *et al*., 2016).

**Fig. 4.**
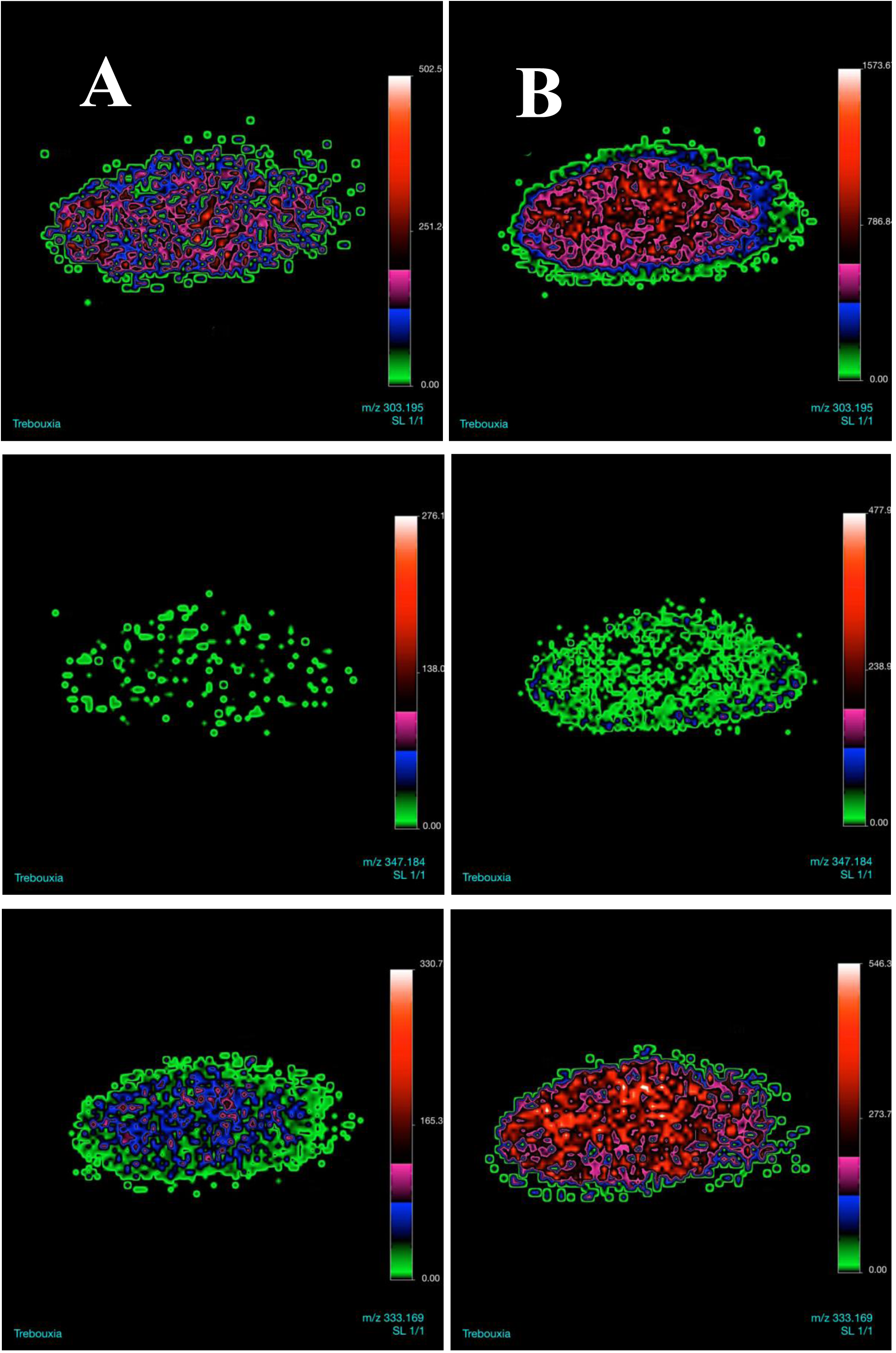
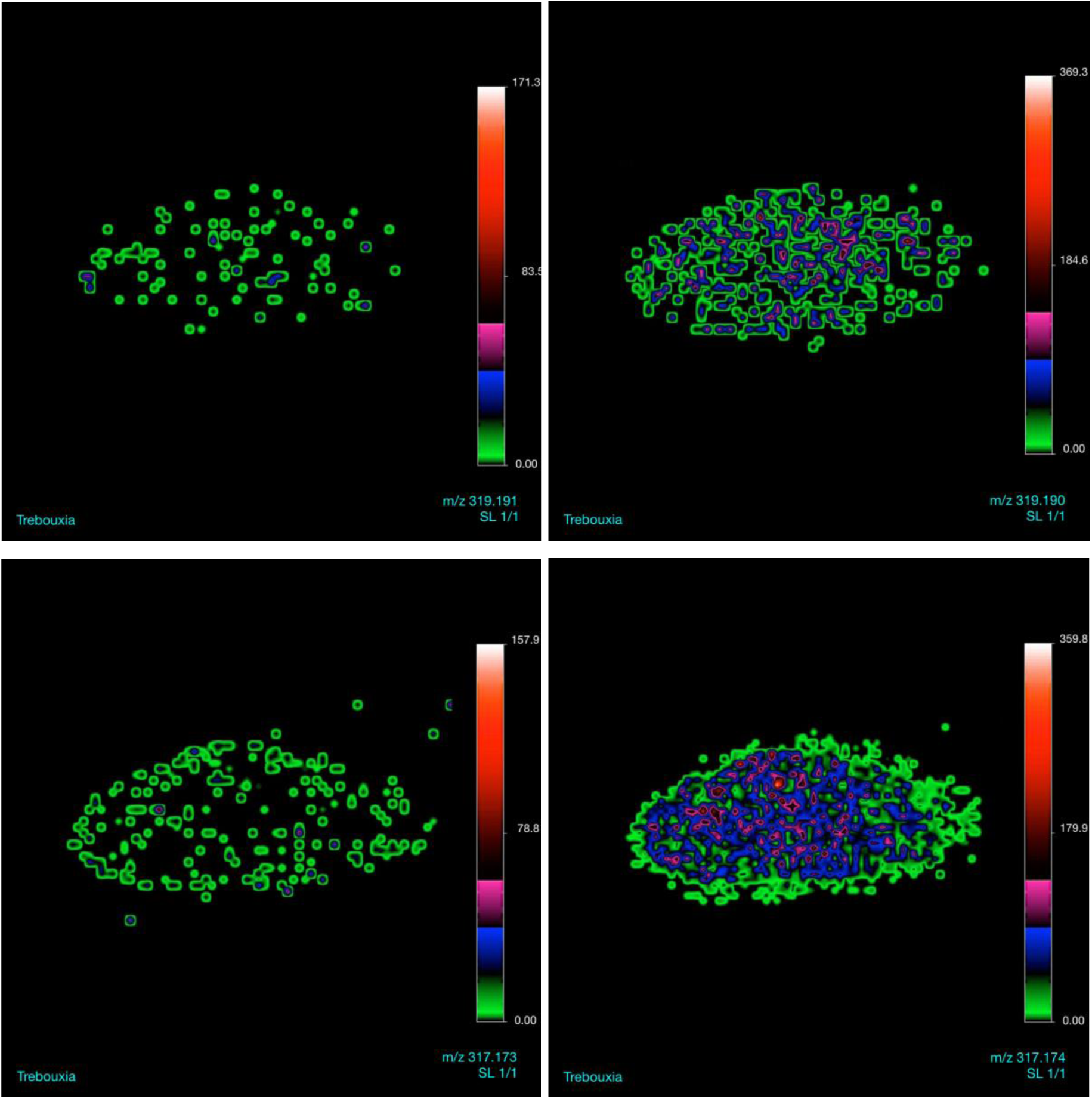
DESI-MSI imaging (positive mode) of carlactones in cells of *Trebouxia* **A**: *Trebouxia* cultivated on media with full strength; **B**: *Trebouxia* was grown on P-free medium before imaging. Images show the localisation of ions: *m/z* 303.195 (carlactone), *m/z* 347.184 (methyl carlactonate), *m/z* 333.169 (carlactonoic acid), *m/z* 319.191 (19-hydroxy carlactone), *m/z* 317.173 (19-oxo carlactone).

### Perspective of practical use of Trebouxia arboricola in pest management

The growth of pea plants cv. Terno, which seeds were treated by algae homogenate, was positively affected by the application, by the arbuscular mycorrhizal fungi or the combined treatment - *Rhizophagus irregularis*, *Trebouxia arboricola* in second short-term experiment. All treatments had stimulation effects on growth characteristics, increase in length of stem and in number of nodes in the most efficient *T. arboricola* plus *R. irregularis*, suggesting that fungi colonization was probably positively affected by the algal treatment (**Table 5**). However the statistical analysis did not show statistically significant differences, especially because of high variability of control. SL levels in pea plant would need to be estimated, and the experiment should be tested in the presence of parasitic seeds (*P. aegyptiaca*), but it can already be concluded that *T. arboricola* application is not phytotoxic and has positive effects on AMF, which is important for the practical purposes.

**Table 5.** Evaluation of growth of pea plants (*Pisum sativum* L., cv. TERNO) after seeds treatment by alga water homogenate or in combination with arbuscular mycorrhizal fungus in pots experiment. Treatments: *Trebouxia arboricola* (*T*); *Rhizophagus irregularis* (*R*). There are not significant differences among the mean values in the table (p > 0.05).

| Treatment | number of nodes | fw of shoots per plant (g) | dw of shoots per plant (g) | length of plant (cm) |
| --- | --- | --- | --- | --- |
| CONTROL | 8.3±1.7 | 1.9±0.7 | 0.215±0.035 | 22.63±6.23 |
| *Trebouxia* | 9.2±0.4 | 2.1±0.5 | 0.218±0.047 | 25.75±2.74 |
| *Rhizophagus* | 8.4±1.4 | 2.0±0.8 | 0.220±0.051 | 25.18±6.54 |
| *R + T* | 8.8±0.4 | 2.0±0.8 | 0.290±0.014 | 25.17±3.88 |

There is accumulating evidence that microalgae (cyanobacteria and algae) like many other plants produce various phytohormones, also including SLs. This fact could be highly interesting from a practical point of view, offering a novel approach for parasitic plant management. Let us assume that *Trebouxia* cells in the artificial cultivation produce and release SL-related compounds into the environment or presence of another strigolactones pathway. There is a vision to implement pilot studies that could verify the use of this biotechnology product in agriculture.

## Supplementary Data

Supplementary data are available at JXB online.

Table S1. DART-HRMS fragmentation analysis (positive mode) of carlactones in fresh samples of *Trebouxia* extract.

Fig. S1. Transcript levels of SL response marker genes PpCCD7 and Pp3c2_34130v3.1 of mutant *Physcomitrella patens Ppccd8* after *Trebouxia* treatment.

## Acknowledgements

The authors would like to thank Catherine Rameau (INRA, AgroParisTech, CNRS, Université Paris-Saclay, Route de St-Cyr 10, 780 26 Versailles Cedex, France), for providing buds outgrowth measurements on rms-1 (*ccd8*) mutant of pea. This study was supported by Grant Agency of MEYS, COST Action FA1206 of Czech Republic, no. LD14101, grant No. GA14-28933S from the Czech Science Foundation; by Scientific Grant Agency of the Ministry of Education of Slovak Republic and the Academy of Sciences (VEGA 2/0138/17); this article is based on work from COST Action FA1206 (STREAM, “STRigolactones Enhances Agricultural Methodologies”), supported by COST (European Cooperation in Science and Technology); by Marie Curie European Re-integration Grant no.239343 and BW/L140-5-0406-0 from the University of Gdańsk.

